# Generation of Unfolded Outer Membrane Protein Ensembles Targeted by Hydrodynamic Properties

**DOI:** 10.1101/2022.10.31.514552

**Authors:** Taylor Devlin, Patrick J. Fleming, Nicole Loza, Karen G. Fleming

## Abstract

Outer membrane proteins (OMPs) must exist as an unfolded ensemble while interacting with a chaperone network in the periplasm of Gram-negative bacteria. Here, we developed a method to model unfolded OMP (uOMP) conformational ensembles using experimental properties of two well-studied OMPs. The overall size and shape of the unfolded ensembles in water were experimentally defined by measuring the sedimentation coefficient as a function of urea concentration. We used these data to model a full range of unfolded conformations by parameterizing a targeted coarse-grained simulation protocol. The ensemble members were further refined by short molecular dynamics simulations to reflect proper torsion angles. The final conformational ensembles reveal inherent differences in the unfolded states that necessitate further investigation. Building these uOMP ensembles advances the understanding of OMP biogenesis and produces essential information for interpreting structures of uOMP-chaperone complexes.

## Introduction

Outer membrane proteins (OMPs) in Gram-negative bacteria encounter several physical challenges to folding. After cytoplasmic translation and translocation across the inner membrane, these unfolded and hydrophobic proteins must cross the periplasm without misfolding or aggregating. While in this aqueous compartment, the unfolded polypeptides encounter a large kinetic barrier to folding into the outer membrane (Barral et al. 2004; Gessmann et al. 2014). To overcome these cellular obstacles, periplasmic chaperones suppress unfolded uOMP aggregation and promote folding competent conformations before uOMPs interact with the β-barrel assembly machine (BAM), which catalyzes their folding into the outer membrane (Hagan et al. 2010; Gessmann et al. 2014; Ulrich and Rapaport 2015; Plummer and Fleming 2016; Chaturvedi and Mahalakshmi 2017; Konovalova et al. 2017; Tomasek and Kahne 2021). These physical challenges and the overall organization of the cell envelope mean that uOMPs must exist in the periplasm in either a free or chaperone-bound state for nontrivial amounts of time (Costello et al. 2016). Insight into the conformations of uOMP ensembles (Krainer et al. 2017) is of great importance for understanding OMP biogenesis in the cell envelope and for building structural models of chaperone-uOMP complexes.

Several methods have been developed to generate and analyze chemically denatured states of classically folded soluble proteins (Fitzkee and Rose 2004; Jha et al. 2005; Curcó et al. 2012) or conformational ensembles of intrinsically disordered proteins (IDPs) (Pelikan et al. 2009; Allison 2017; Bonomi et al. 2017; Shrestha et al. 2019; Ahmed et al. 2020; Larsen et al. 2020; Tesei et al. 2021). Coarse-grained and all-atom simulations utilizing various force fields, simulation conditions, and intra-molecular restraints have been the computational foundation of such methods. The primary goal of these methods is to create a structural ensemble described by calculated properties in agreement with experimental properties (Bernadó et al. 2007; Róycki et al. 2011; Antonov et al. 2016; Shevchuk and Hub 2017; Potrzebowski et al. 2018; Shrestha et al. 2019; Bottaro et al. 2020; Ahmed et al. 2021; Tesei et al. 2021). Experimental properties indicative of the overall size and shape of a collection of unfolded, denatured, or disordered polypeptides include the radius of gyration and maximum dimension determined by scattering methods; rotational diffusion determined by nuclear magnetic resonance (NMR) or fluorescence methods; and translational diffusion determined by analytical ultracentrifugation (AUC), NMR or single molecule fluorescence methods. To match computational and experimental values, simulation conditions are configured to bias the resulting ensemble toward the experimental value directly, or alternately, the initial unbiased ensemble is trimmed or weighted to provide agreement with experimental data.

We built upon the ideas and procedures described above to develop a simulation procedure that creates uOMP ensembles consistent with experimentally determined hydrodynamic properties. The procedure described here uses a two-step protocol with a coarse-grained molecular dynamics (MD) first step to create a well-sampled conformational ensemble followed by an all-atom MD second step to relax the stereochemistry of the polypeptide.

Unlike typical soluble proteins or IDPs, OMPs contain several hydrophobic segments corresponding to transmembrane strands. Therefore, we reason that hydrophobic interactions play a significant role in dictating the structural properties of the uOMP ensemble in aqueous solutions. To control the degree of hydrophobic collapse during simulations, we used the coarse-grained molecular dynamics simulation software application CafeMol (Kenzaki et al. 2011) for our initial simulations of the unfolded state. CafeMol has an easily configured force field with a tunable hydrophobic potential term and does not require solvent atoms. To experimentally characterize the average size and shape of an uOMP, we used sedimentation velocity analytical ultracentrifugation (SV-AUC) to determine sedimentation coefficients as a function of chemical denaturant. Although scattering methods are most frequently employed for soluble proteins (Riback et al. 2017; Bowman et al. 2020; Ahmed et al. 2021), the high concentrations required for those experiments are not accessible to unfolded membrane proteins. In contrast, sedimentation velocity can be conducted at much lower protein concentrations below the threshold for aggregation. Comparison of experimental and calculated sedimentation coefficients for the thousands of models generated during a simulation requires a rapid method to calculate hydrodynamic properties. HullRad is a fast and accurate program that works with both folded and unfolded protein structural models (Fleming and Fleming 2018). Here we describe a procedure to generate ensembles of uOMPs with the coarse-grained MD engine CafeMol followed by all-atom MD, and we use the program HullRad to calculate hydrodynamic properties of generated model ensembles for comparison to experimental sedimentation coefficients. The final ensembles of two uOMPs reveal variation in the unfolded state properties attributable to amino acid composition or sequence differences that warrant further investigation.

## Materials and Methods

### Sedimentation Velocity as a Function of Urea Concentration

Both unfolded OmpA_171_ (uOmpA_171_) and unfolded OmpX (uOmpX) were diluted to 2 μM in 1, 2, 4, 6, or 8 M urea with either a 20 mM Tris or 20 mM sodium phosphate, pH 8 background buffer for SV-AUC. Samples were prepared and centrifuged in triplicate. All SV-AUC experiments were performed using a Beckman XL-A ultracentrifuge (Beckman Coulter) and cells with 1.2 mm double-sector epoxy centerpieces and sapphire windows. Each sample was centrifuged at 25 °C using a 4-hole, An-60Ti rotor at a rotor speed of 50,000 rpm. Radial scans at 230 nm were acquired with 0.003 cm radial steps in continuous mode with zero time interval between scans. Prior to starting each run, the rotor was temperature equilibrated in the instrument for at least 60 minutes.

All SV-AUC data sets were analyzed using dc/dt+ (Philo 2006). Sedimentation coefficient distributions (g(s*) distributions) were corrected to 20 °C in water using the appropriate densities (ρ), viscosities (η), and partial specific volumes 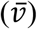 for each buffer and protein calculated using SEDNTERP. These values are presented in Supplemental Table 1, which also shows the buffer-corrected experimental average (<s_20,w_>), corrected fit (s_20,w_) sedimentation coefficients, and the experimentally determined molar mass values. Plots of s_20,w_ versus urea concentration were fit to a line (y = mx + y_0_) to extrapolate the sedimentation coefficient of the uOMP in 0 M urea (s_0_) from the y-intercept. Errors reported on the y-intercept represent the 95% confidence intervals from globally fitting all sedimentation coefficients collected in triplicate for each protein at each urea concentration. The f/f_0_ and axial ratio for uOmpA_171_ in water were calculated in SEDNTERP using the Teller method and the default hydration value.

**Table 1.** Parameters from linearly fitting plots of s_20,w_ versus urea concentration. Three replicates were performed at each urea concentration, and all data was globally fit to a linear equation. Errors are reported as the 95% confidence interval.

| **Protein** | **Buffer Background** | **Slope**<br>Svedbergs/M | **Y-Intercept (95% CI)**<br>Svedbergs |
| --- | --- | --- | --- |
| uOmpA<sub>171</sub> | 20 mM sodium phosphate | -0.040 | 1.59 (1.58-1.60) |
|  | 20 mM tris | -0.036 | 1.61 (1.59-1.63) |
| uOmpX | 20 mM sodium phosphate | -0.038 | 1.53 (1.52-1.54) |
|  | 20 mM tris | -0.034 | 1.53 (1.52-1.54) |

### Generation of Structural Ensembles

For each protein a heavy atom model was built from the UNIPROT sequence of the protein using PyMOL (PyMol Molecular Graphics System, Version 2.5.0, Schrodinger, LLC). An extended conformation was obtained using backbone torsion angles ϕ=−75° and ψ=145°. We performed coarse-grained MD simulations on the extended protein model using CafeMol (Kenzaki et al. 2011), which first converts the protein chain into a random C_α_-only chain conformation. Simulations were run at 298 K using Langevin dynamics with residue-specific mass, a flexible local potential, excluded volume repulsive interaction, and a hydrophobic interaction potential. Step size was 0.4, and total simulation steps equaled 2.5 x 10^6^ with a conformation saved every 10,000 steps. A series of different simulations for each protein were run at different coefficients of hydrophobic interaction (described below), and a replicate simulation was run on each protein at the optimal hydrophobic interaction coefficient to ensure consistency of results.

Every other frame from the last 2000 frames of the coarse-grained simulation was saved using CATDCD (Humphrey et al. 1996) to ensure non-correlated sampling of the trajectory. VMD (Humphrey et al. 1996) was used to write 1000 PDB files of C_α_-only structures, and PULCHRA (Rotkiewicz and Skolnick 2008) was used to rebuild side chains and back map the amino acid residue. All-atom MD simulations in implicit solvent were carried out on each of the 1000 structures using NAMD (Phillips et al. 2005) to relax van der Waals (VDW) clash and obtain Ramachandran compatible backbone torsion angles. These simulations were controlled with CHARMM 36 (Huang and Mackerell 2013) at 298 K with Langevin temperature control under generalized Born implicit solvent conditions, ion concentration = 0.3, and α cutoff = 12.0. The system was minimized with 1000 steps, and MD continued for 25000 steps with a 1.0 fs time step. Hydrodynamic properties were calculated using HullRad (Fleming and Fleming 2018) for the ensemble of structures.

All computer methods described here may be carried out with Mac OS or LINUX machines; the software is freely available.

## Results

### Urea-Dependent Sedimentation Coefficients Determined by SV-AUC

To simulate unfolded outer membrane protein ensembles, experimentally derived structural properties are required. Scattering techniques have traditionally been employed for this purpose, however the high concentrations needed for scattering are problematic for unfolded outer membrane proteins because they aggregate and are no longer monomeric (Tan et al. 2010). An alternative experimental parameter that can be determined at low protein concentrations is the sedimentation coefficient. We therefore performed sedimentation velocity on uOmpA_171_ and uOmpX in several urea concentrations between 1 and 8 M. Both uOMPs were found to be monomeric and monodisperse at protein concentrations of 2 μM in urea concentrations as low as 1 M. Figures 1A and B show the g(s*) distributions of uOmpA_171_ and uOmpX respectively at five different urea concentrations. The proteins both sediment and diffuse more slowly at higher urea concentrations due partly to the increased density and viscosity of the solvent. However, even after correcting for the buffer density and viscosity (i.e. correcting to s_20,w_), the sedimentation of the uOMP still depends on the urea concentration indicating that urea also affects the overall expansion or compaction of the uOMP conformational ensemble (Fig. 1C and D). While experiments in urea concentrations lower than 1 M are inaccessible due to protein aggregation, a sedimentation coefficient in 0 M urea (s_0_) most accurately describes the possible uOMP ensemble in an aqueous environment. We can estimate this value, termed s_0_, by extrapolating the s_20,w_ values as a function of the urea concentration. Both uOmpA_171_ and uOmpX show linear extrapolations to 0 M urea (Fig. 1C and D). Sedimentation coefficients decrease with increasing urea concentration (negative slope) for both uOMPs, indicating that uOMP ensembles in high urea concentrations are more expanded than ensembles in low urea concentrations. We were curious about the effect of buffer on our measurements, so we repeated all SV-AUC experiments in both a phosphate and Tris buffer. The buffer background does not impact the overall compaction of the ensemble at 0 M urea, and data collected for uOmpA_171_ in a Tris background agrees with previously published results (Supplemental Fig. 1) (Danoff and Fleming 2011). All data, fit parameters, and errors for each dataset are listed in Table 1 and Supplemental Table 1. The measured s_0_ for uOmpA_171_ = 1.59 and uOmpX = 1.53 Svedbergs were used to target our simulated uOMP ensembles.

**Fig 1.**
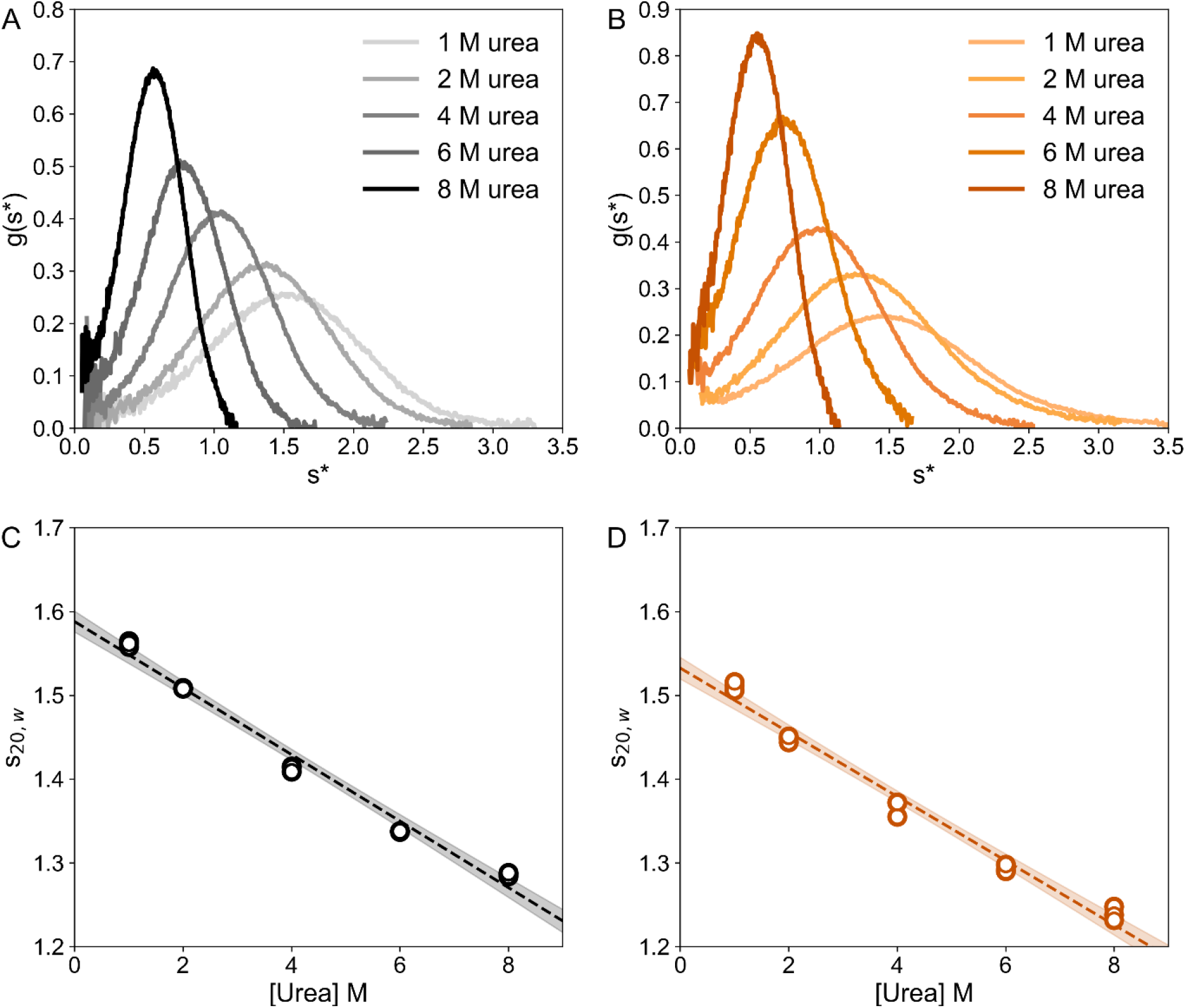
The sedimentation of uOmpA_171_ and uOmpX depends linearly on the urea concentration. (A) Representative g(s*) distributions of uOmpA_171_ in 1, 2, 4, 6, and 8 M urea. (B) Representative g(s*) distributions of uOmpX in 1, 2, 4, 6, and 8 M urea. (C) uOmpA_171_ s_20,w_ versus urea concentration fits to a line with the equation y = −0.04x + 1.59. (D) uOmpX s_20,w_ versus concentration fits to a line with the equation y = −0.04x + 1.53. For both uOMPs, the protein concentration was 2 μM in a buffer of 20 mM sodium phosphate plus the experimental urea concentration between 1 and 8 M, pH 8. Absorbance was measured at 230 nm while spinning at 50000 rpm and 25 °C. Data were analyzed using dc/dt+ to determine g(s*) distributions and s_20,w_ values. Each experiment was performed in triplicate, and all three data points at each urea concentration are included in the liner fit. Shaded regions represent the 95% confidence interval on the fit line.

### Generation of Conformational Ensembles: Tuning the Hydrophobic Interaction Potential

Unfolded ensembles of the two model OMPs were created as described in the methods. using coarse-grained MD simulations targeted to the experimentally determined s_0_. Extended, heavy atom protein models were built in PyMOL and converted to C_α_-only bead models for simulation in the coarse-grained MD engine CafeMol. The complete force field used during CafeMol simulations consists of four pseudo-energy terms: 1) volume exclusion; 2) backbone angle; 3) backbone torsion; and 4) hydrophobic potential. The backbone angles, backbone torsions, and hydrophobic potentials are amino acid specific, and default values were used. The hydrophobic interaction potential is defined as,

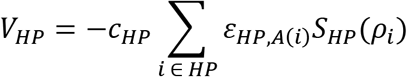

where *c*_HP_ scales the overall strength of the hydrophobic interactions, *ε*_HP,A(i)_ is a residue-specific hydrophobicity factor, and *S*_HP(_ρ_i)_ quantifies the degree of “buriedness” of the residue (Kenzaki et al. 2011). Here we use the *c*_HP_ term to bias the compaction of unfolded OmpA_171_ and OmpX during coarse-grained simulation by tuning the strength of the hydrophobic interaction potential.

We systematically incremented *c*_HP_ between 0.5 and 1.2 until the compaction of our simulated ensembles reflected the experimentally determined values of s_0_. For the selection of an appropriate *c*_HP_, 1000 coarse-grained models generated by CafeMol were converted to all-atom models using PULCHRA. The degree of compaction was quantified using the ensemble-average sedimentation coefficient calculated using HullRad. Fig. 2 shows the effect of varying *c*_HP_ on the calculated average sedimentation coefficients for both uOmpA_171_ and uOmpX.

**Fig 2.**
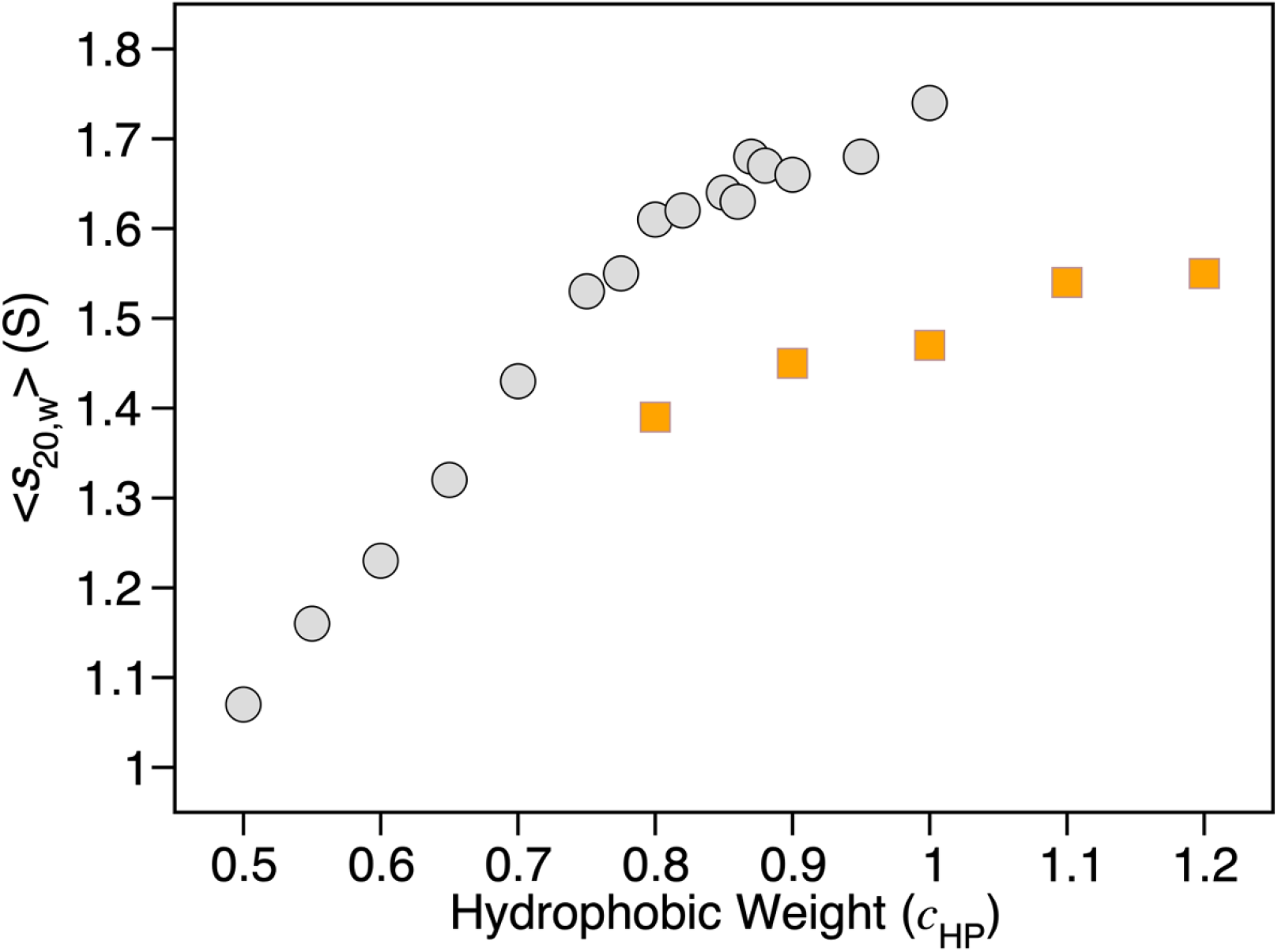
The effect of the hydrophobic coefficient on the CafeMol modeled sedimentation coefficient. The value of *c*_HP_ during coarse-grained simulation with CafeMol varied between 0.5 and 1.2, and the average sedimentation coefficients were calculated from each ensemble of 1000 structures using HullRad. Grey data points, uOmpA_171_; orange data points, uOmpX.

For uOmpA_171_, *c*_HP_=0.87 results in an initial ensemble average sedimentation coefficient (<s_20,w_>) of 1.62 Svedbergs. Further refinement of the ensemble as described below slightly expands the structures to result in an ensemble average of 1.59 Svedbergs for uOmpA_171_, in agreement with the experimentally determined value. For uOmpX the corresponding simulation *c*_HP_ is 1.1, which results in an initial ensemble <s_20,w_> of 1.54 Svedbergs before and 1.50 Svedbergs after refinement as described below.

### Generation of Conformational Ensembles: Further Structural Refinement

Amino acid side chains were rebuilt on the CafeMol coarse-grained C_α_-only structures using PULCHRA as described in the Methods. However, this results in some atomic van der Waals (VDW) clash as well as unfavorable backbone torsion angles. These unfavorable backbone dihedral angles are shown for all alanine residues in the 1000-member ensemble of uOmpA_171_ in Fig. 3 (green dots). We used further structural refinement to minimize these unfavorable energy states in the final conformational ensembles. Short all-atom molecular dynamics simulations in implicit solvent restore favorable backbone torsion angles (Fig. 3, black dots). This is true not only for alanine but for all residue types found in either OmpA_171_ or OmpX. After the short all-atom simulations, the Ramachandran plots of all residue types agree qualitatively with backbone phi (ϕ) and psi (ψ) values from an unbiased coil library (Beck et al. 2008) (Supplemental Fig. 2).

**Fig 3.**
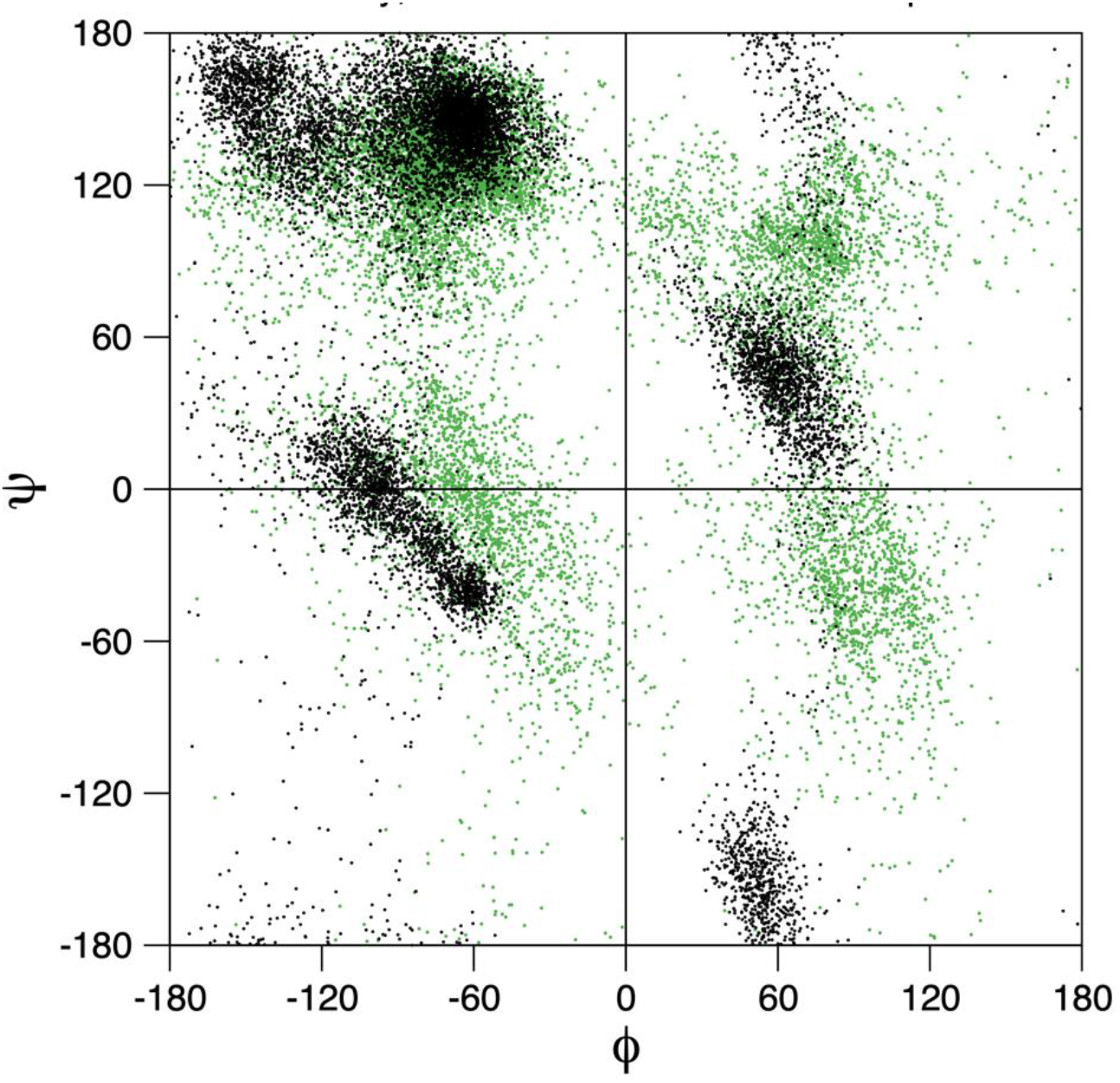
Non-favorable backbone dihedral angles in initial back mapped structures are relieved by all-atom simulation. Ramachandran plot of ALA backbone angles in a 1000 member uOMP ensemble before (green) and after (black) refinement using a short all-atom MD simulation. The black data are consistent with the ALA ϕ, ψ angle distribution observed in an unbiased coil library (Beck et al. 2008).

The two-part protocol described here, including both a coarse-grained and all-atom simulation, was designed to ensure extensive sampling of available conformational space of the proteins. Even within the confines of allowed ϕ and ψ angles determined by the coarse-grained force field, the refined models still exhibit a wide range of backbone conformations with an average deviation of greater than 50° across the whole protein (Fig. 4 A and B). Proline residues are the exception with significantly smaller ϕ deviations.

**Fig 4.**
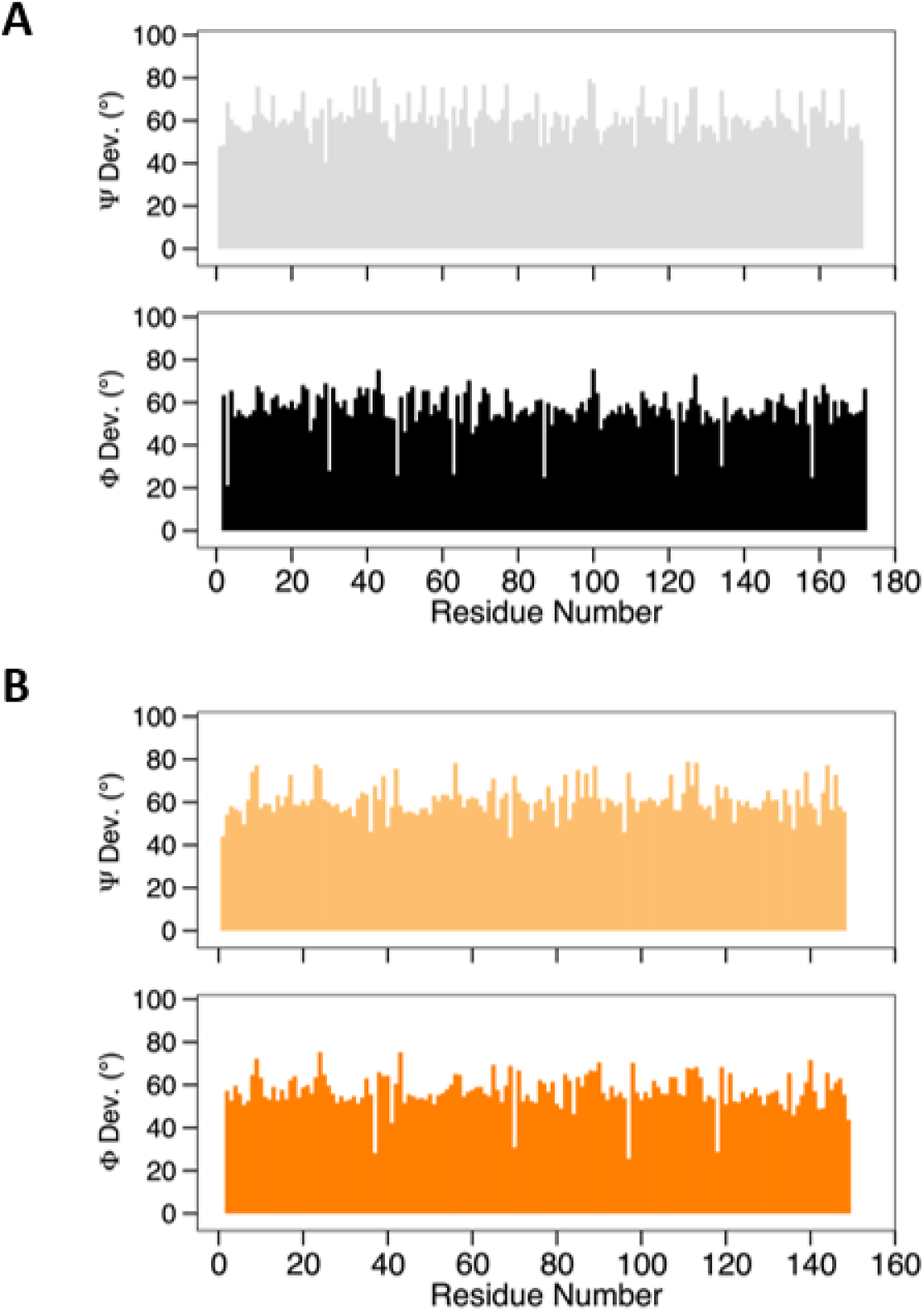
Refined all-atom simulation models exhibit a wide range of backbone conformations. The average angular standard deviations of backbone ψ and ϕ angles of 1000 refined structures for (A) uOmpA171 and (B) uOmpX are plotted as bar graphs. The very low ϕ angular deviations are observed for proline residues.

### Final Ensembles Cover a Wide Range of Unfolded States

Coarse-grained unfolded ensembles were generated with the correctly tuned hydrophobic potential, converted to all-atom models, and refined to alleviate VDW clashes and fix unfavorable torsion angles. The final ensembles contain approximately 1000 conformations (occasionally VDW overlaps were not relieved during minimization and these simulations were discarded). We used HullRad to quickly calculate the ensemble average hydrodynamic properties and the hydrodynamic properties of each individual conformation. The distributions of calculated sedimentation coefficients (s_20,w_) for uOmpA_171_ and uOmpX are shown in Fig. 5. uOmpA_171_ has a non-symmetrical distribution skewed toward more expanded conformations with smaller sedimentation coefficients, whereas uOmpX exhibits an approximately normal distribution. Despite having the same number of β-strands and similar molecular weights, these two uOMPs present different distributions, indicating that properties of the ensemble derive from amino acid composition or sequence. Duplicate simulations confirm that sequence specific differences in the distributions of calculated sedimentation coefficients are reproducible (data not shown).

**Fig 5.**
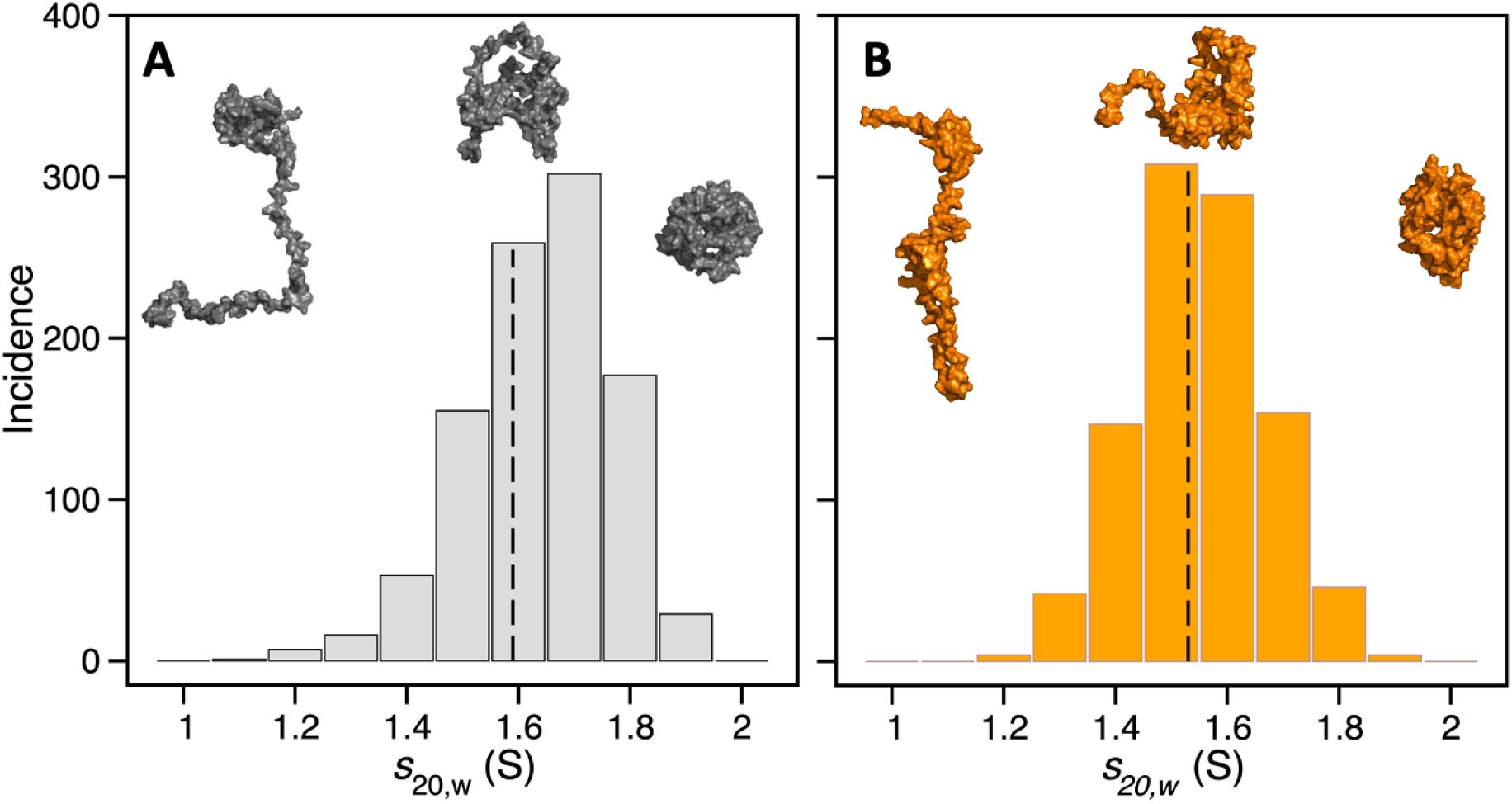
Refined model calculated sedimentation coefficients of (A) uOmpA_171_ and (B) uOmpX show wide distributions. Three representative atomic models from across the distribution are shown for each uOMP. The average calculated sedimentation coefficient from the distribution of uOmpA_171_ is 1.59 S; a vertical dashed line marks the experimental sedimentation coefficient of 1.59 S. The average calculated sedimentation coefficient from the distribution of uOmpX is 1.50 S; a vertical dashed line marks the experimental sedimentation coefficient of 1.53 S.

All models of the same protein have the same molar mass, which means that the spread of the calculated sedimentation coefficients reflects a wide range of expansion or contraction represented in the final ensembles. Compact, molten globule-like, and extended conformations are found in each ensemble. Examples of conformations across the full range of the distribution are shown as atomic sphere models in Fig. 5. Additional evidence for a wide range of conformational states in the uOMP ensembles is shown in histograms of the N-terminal to C-terminal distance ranging from ~10 Å to >100 Å (Fig. 6). The end-to-end distance distributions are consistent with the sedimentation coefficient distributions in Fig. 5 in that the uOmpA_171_ distribution is skewed towards more expanded structures while the uOmpX distribution is approximately normal. The difference in expansion (or contraction) is also reflected in the mean end-to-end distance with the uOmpX ensemble having a mean distance of 61.9 Å and the uOmpA_171_ ensemble having a mean distance of 39.6 Å.

**Fig 6.**
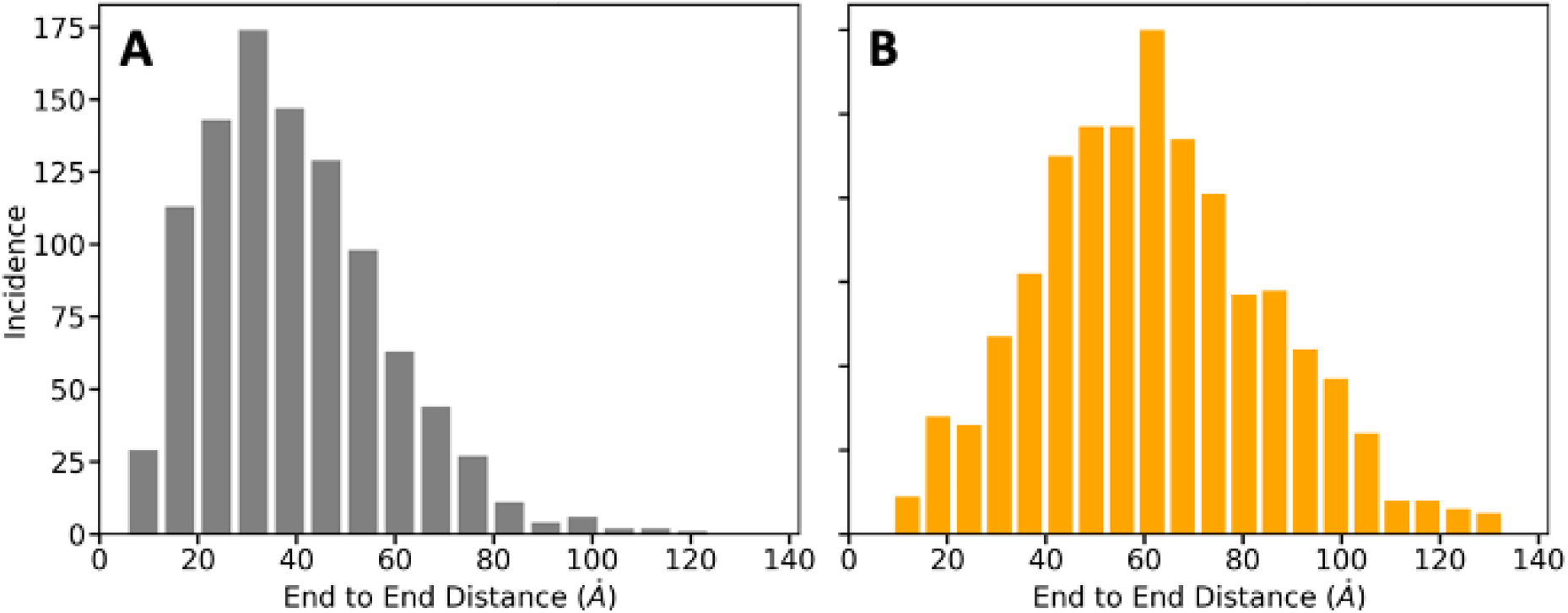
Model ensembles of (A) uOmpA_171_ and (B) uOmpX contain different distributions of end-to-end distances. Distances between the C_α_ atoms of the first and last residues of each uOMP model were calculated and plotted as histograms. The uOmpA_171_ mean end-to-end distance = 39.6 (SD 18.7) Å, and for uOmpX the mean end-to-end distance = 61.9 (SD 23.3) Å.

The experimental s_0_ for uOmpA_171_ in phosphate buffer is 1.59 (1.58-1.60). Using this value we calculated a frictional ratio (f/f_o_) of 1.61 in SEDNTERP. The frictional ratio indicates the additional frictional drag on a macromolecule due to hydration and shape asymmetry over and above that of a sphere of equivalent molecular mass. Assuming a hydration of 0.387 g/g (calculated from sequence in SEDNTERP), this frictional ratio would indicate an average axial ratio ensemble of 7.6:1 for the uOmpA_171_ ensemble, a value consistent with previously reported results (Danoff and Fleming 2011). In contrast, HullRad calculates an average axial ratio of 2.0:1 for the uOmpA_171_ ensemble, and this degree of asymmetry is more consistent with the distributions of conformations observed in the movies illustrating sampling of the uOMP ensembles (Supplementary Movies 1 and 2). This more reasonable value for the axial ratio would suggest that uOMPs have more than 0.387 g/g of hydration. Using the methods described elsewhere in this issue, it should now be possible to investigate the hydration levels of uOMPs and other unfolded or disordered proteins (Fleming et al. 2022).

## Discussion

During OMP biogenesis in Gram-negative bacteria, unfolded OMPs traverse the aqueous periplasm before folding into the outer membrane. Periplasmic chaperones bind uOMPs during this process to prevent aggregation and further facilitate folding. To fully understand the folding pathways of these bacterial membrane proteins, it is important to delineate the conformational states of free uOMPs and determine how binding to periplasmic chaperones alters these states. Knowledge about uOMP ensembles may even explain how the nascent proteins are recognized by periplasmic proteins in the first place. Additionally, access to uOMP ensembles is required to build structural models of a chaperone-uOMP complex, as has recently been done for SurA binding uOmpA (Marx et al. 2020). Here we describe a protocol for generating structural ensembles of uOMPs that reflect the experimentally determined properties of size and shape.

We used SV-AUC to measure the sedimentation coefficient of two different uOMPs, uOmpA_171_ and uOmpX, as a function of urea concentration. The negative slope of the uOMP sedimentation coefficient as a function of urea concentration indicates that the unfolded ensembles are more expanded in higher urea concentrations and more collapsed in lower urea concentrations. These results are consistent with the expansion of unfolded, disordered, or denatured ensembles in high concentrations of chemical denaturant reported in the literature (Sherman and Haran 2006; Tezuka-Kawakami et al. 2006; Hofmann et al. 2012; Aznauryan et al. 2016; Zheng et al. 2016; Peran et al. 2019). We extrapolated the data to 0 M urea to determine the sedimentation coefficient of the aqueous ensemble (s_0_) and used this property to target simulations. While experiments in the absence of urea are inaccessible due to aggregation, extrapolation to 0 M urea is an essential step to determine the aqueous properties of the unfolded ensemble in an environment like the periplasm because there is a distinction between the physiologically relevant, aqueous unfolded state and the chemically denatured state. The urea-dependence of the uOMPs global hydrodynamic properties like their sedimentation coefficient serves as a reminder that the presence of even low urea concentrations may affect the properties of uOMPs.

We developed a protocol to build and target unfolded protein ensembles using the experimentally determined s_0_ value. The procedure described here for generating unfolded ensembles is similar to that described by Curcó et al. in that it proceeds with two steps: 1) Creating a set of structures that explore a large range of conformations within favorable backbone dihedrals, and 2) Relaxing and refining the initial set of structures using an all-atom force field. In the procedure described here molecular dynamics is used for both the initial and refinement stage in contrast to the Monte Carlo algorithm used by Curcó et al (Curcó et al. 2012). The hydrophobic potential term, *c*_HP_, controls the hydrophobic interaction of amino acids during coarse-grained simulations and is the term used to tune overall hydrophobic collapse in the initial simulation. In this way the resulting structural ensembles do not require subsequent filtering or weighting to obtain the appropriate ensemble average sedimentation coefficient.

Although a primary goal of ensemble generation is to match calculated hydrodynamic properties with experimental values, another important objective is to obtain a reasonable sampling of all possible conformational states in the ensemble. The ensembles generated by the protocol described here exhibit a wide range of backbone conformations, degrees of compaction or expansion, and differences in end-to-end distances. Additional experimental data on intrinsic structure such as specific amino acid residue distances or backbone dihedral angles would be useful in further validating these types of ensembles.

The unfolded ensembles obtained for each uOMP represent a range of expanded, molten globule-like, and collapsed conformations (Fig. 5). However, the generated ensembles of the two uOMPs produce distinct distributions; while conformations of uOmpX are normally distributed, the uOmpA_171_ distribution is skewed toward smaller sedimentation coefficients and therefore more expanded conformations. A different hydrophobic coefficient was required to achieve the correct ensemble average sedimentation coefficient for each uOMP. These differences imply that the unfolded ensembles of these two similarly sized OMPs depends on amino acid composition or sequence. Therefore, it would be interesting to explore how this method translates to modeling the unfolded states of several larger uOMPs. A more extensive survey of uOMPs would also reveal any length dependent trends in s_0_ or hydrophobic potential and may explore the sequence determinants of unfolded ensemble properties including the effects of global hydrophobic content, local clustering of hydrophobic residues, and the role of charged residues (Bowman et al. 2020).

Riback et al. show that water is a “good solvent” for an unfolded but foldable soluble protein, resulting in an expanded denatured state ensemble (Riback et al. 2017). This data runs counter to widespread ideas that hydrophobic interactions drive collapse of unfolded ensembles in water. Here use of a hydrophobic interaction potential in the coarse-grained simulations is required to compact the initial extended chain models until they match experimentally-derived sedimentation coefficients. This implies that some degree of hydrophobic collapse occurs in these systems. However, the resulting ensembles contain both expanded and collapsed conformations (Fig. 5 and 6, Supplemental Movies 1 and 2). To compare the state of collapse of the uOMPs described here with other unfolded state ensembles we fit the average R_G_ values to the Flory function R_G_ = R_0_N^*ν*^ with R_0_ set to 2.0 (Fitzkee and Rose 2004). In contrast to the results shown by Riback et al., the average R_G_ for both uOmpA_171_ and uOmpX fit to the function with a scaling factor (*ν*) of 0.5 (Fig. 7). This result indicates that the uOMPs are more collapsed than an unfolded soluble protein with a scaling factor of >0.5. To our knowledge, this work marks the first to comprehensively model the unfolded ensemble of a membrane protein in aqueous solution. We acknowledge that the sample size presented in this paper is small, but these initial observations are intriguing, and it will be interesting to determine whether these observations will hold true for a range of different uOMP ensembles. Future investigations of uOMPs of varying sequence and molar mass will address this question.

**Fig 7.**
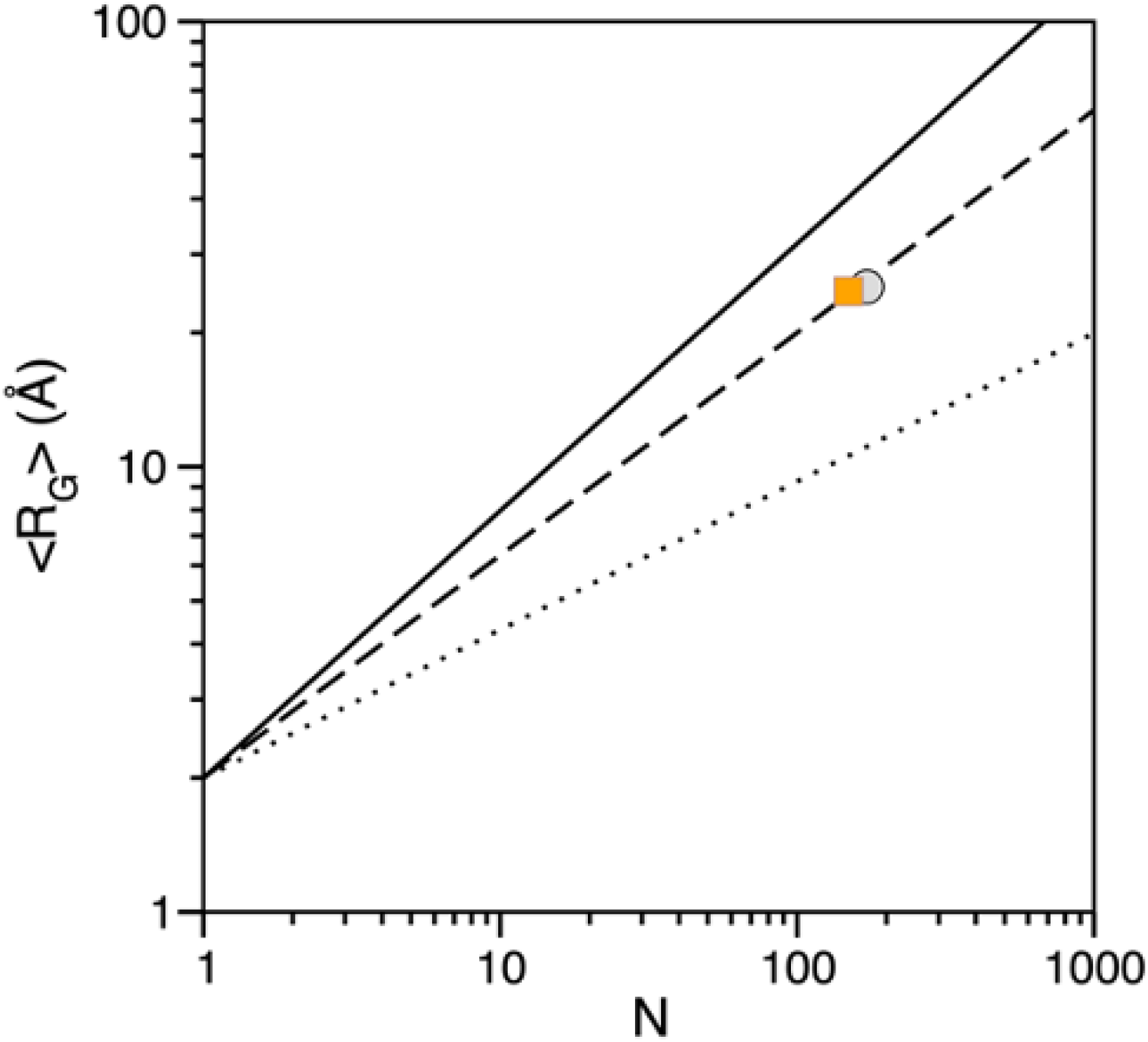
The calculated radius of gyration (R_G_) for uOMPs reflects ideal random-coil behavior in a good solvent. The ensemble average R_G_ for uOmpA_171_ (grey circle) and uOmpX (orange square) are plotted versus number of amino acid residues. The lines reflect the function R_G_=2.0N^*ν*^ where *ν* equals 0.6 (solid line), 0.5 (dashed line), 0.33 (dotted line).

The use of coarse-grained simulations for the initial ensembles and HullRad to connect computational models and experimental data makes for a simple, modifiable, and time-efficient method to generate unfolded ensembles. Here we use the sedimentation coefficient to direct ensemble generation, but other hydrodynamic properties such as radius of gyration or translational diffusion coefficient may also be easily implemented in this procedure because HullRad calculates a full suite of hydrodynamic and physical properties. The procedure can also be extended to other unfolded but foldable protein systems such as unfolded membrane proteins in the mitochondrial inner membrane space due to the tunable hydrophobic potential term in CafeMol. The computationally-created ensembles of uOMPs allow for analysis of intrinsic properties of the unfolded state, for exploring the mechanism of recognition by periplasmic chaperones, and for building structural models of chaperone-uOMP complexes.

## Supplementary Information

The online version contains supplementary material available in OMPEnsembles_Supplement.pdf. Supplementary movies are submitted as OmpA171_Movie.mp4 and OmpX_Movie.mp4.

## Authors contributions

TD, PJF and KGF conceived the experimental project. KGF directed the research. TD and NL carried out laboratory experiments and analyzed the data; PJF conducted MD simulations and computational analysis; TD, PJF and KGF wrote the manuscript. All authors read and approved the final manuscript.

## Availability of data and materials

All data will be freely available upon request.

## Acknowledgments

This work was carried out at the Advanced Research Computing at Hopkins (ARCH) core facility (rockfish.jhu.edu), which is supported by the National Science Foundation (NSF) grant number OAC1920103.We thank members of the Fleming Lab for helpful feedback.

## References

Ahmed MC, Crehuet R, Lindorff-Larsen K (2020) Computing, Analyzing, and Comparing the Radius of Gyration and Hydrodynamic Radius in Conformational Ensembles of Intrinsically Disordered Proteins. In: Kragelund BB, Skriver K (eds) Intrinsically Disordered Proteins: Methods and Protocols. Springer Science+Business Media, LLC, pp 429–445

Ahmed MC, Skaanning LK, Jussupow A, et al (2021) Refinement of α-Synuclein Ensembles Against SAXS Data: Comparison of Force Fields and Methods. Front Mol Biosci 8:1–13. https://doi.org/10.3389/fmolb.2021.654333

Allison JR (2017) Using simulation to interpret experimental data in terms of protein conformational ensembles. Curr Opin Struct Biol 43:79–87. https://doi.org/10.1016/j.sbi.2016.11.018

Antonov LD, Olsson S, Boomsma W, Hamelryck T (2016) Bayesian inference of protein ensembles from SAXS data. Phys Chem Chem Phys 18:5832–5838. https://doi.org/10.1039/c5cp04886a

Aznauryan M, Delgado L, Soranno A, et al (2016) Comprehensive structural and dynamical view of an unfolded protein from the combination of FRET, NMR, and SAXS. Proc Natl Acad Sci U S A E5389–E5398. https://doi.org/10.1073/pnas.1607193113

Barral JM, Broadley SA, Schaffar G, Hartl FU (2004) Roles of molecular chaperones in protein misfolding diseases. Semin. Cell Dev. Biol. 15:17–29

Beck DAC, Alonso DOV, Inoyama D, Daggett V (2008) The intrinsic conformational propensities of the 20 naturally occurring amino acids and reflection of these propensities in proteins. Proc Natl Acad Sci U S A 105:12259–12264. https://doi.org/10.1073/pnas.0706527105

Bernadó P, Mylonas E, Petoukhov M V., et al (2007) Structural characterization of flexible proteins using small-angle X-ray scattering. J Am Chem Soc 129:5656–5664. https://doi.org/10.1021/ja069124n

Bonomi M, Heller GT, Camilloni C, Vendruscolo M (2017) Principles of protein structural ensemble determination. Curr Opin Struct Biol 42:106–116. https://doi.org/10.1016/j.sbi.2016.12.004

Bottaro S, Bengtsen T, Lindorff-Larsen K (2020) Integrating Molecular Simulation and Experimental Data: A Bayesian/Maximum Entropy Reweighting Approach. Methods Mol Biol 2112:219–240. https://doi.org/10.1007/978-1-0716-0270-6_15

Bowman MA, Riback JA, Rodriguez A, et al (2020) Properties of protein unfolded states suggest broad selection for expanded conformational ensembles. Proc Natl Acad Sci U S A 117:23356–23364. https://doi.org/10.1073/pnas.2003773117

Chaturvedi D, Mahalakshmi R (2017) Transmembrane β-barrels: Evolution, folding and energetics. Biochim Biophys Acta - Biomembr 1859:2467–2482. https://doi.org/10.1016/j.bbamem.2017.09.020

Costello SM, Plummer AM, Fleming PJ, Fleming KG (2016) Dynamic periplasmic chaperone reservoir facilitates biogenesis of outer membrane proteins. Proc Natl Acad Sci U S A 113:4794–4800. https://doi.org/10.1073/pnas.1601002113

CurcÓ D, Michaux C, Roussel G, et al (2012) Stochastic simulation of structural properties of natively unfolded and denatured proteins. J Mol Model 18:4503–4516. https://doi.org/10.1007/s00894-012-1456-6

Danoff EJ, Fleming KG (2011) The soluble, periplasmic domain of OmpA folds as an independent unit and displays chaperone activity by reducing the self-association propensity of the unfolded OmpA transmembrane β-barrel. Biophys Chem 159:194–204. https://doi.org/10.1016/j.bpc.2011.06.013

Fitzkee NC, Rose GD (2004) Reassessing random-coil statistics in unfolded proteins. Proc Natl Acad Sci U S A 101:12497–12502. https://doi.org/10.1073/pnas.0404236101

Fleming PJ, Correia JJ, Fleming KG (2022) Revisiting Macromolecular Hydration with HullRadSAS. bioRxiv. https://doi.org/https://doi.org/10.1101/2022.10.20.513022

Fleming PJ, Fleming KG (2018) HullRad: Fast Calculations of Folded and Disordered Protein and Nucleic Acid Hydrodynamic Properties. Biophys J 114:856–869. https://doi.org/10.1016/j.bpj.2018.01.002

Gessmann D, Chung YH, Danoff EJ, et al (2014) Outer membrane β-barrel protein folding is physically controlled by periplasmic lipid head groups and BamA. Proc Natl Acad Sci U S A 111:5878–5883. https://doi.org/10.1073/pnas.1322473111

Hagan CL, Kim S, Kahne D (2010) Reconstitution of outer membrane protein assembly from purified components. Science (80-) 328:890–892. https://doi.org/10.1126/science.1188919

Hofmann H, Soranno A, Borgia A, et al (2012) Polymer scaling laws of unfolded and intrinsically disordered proteins quantified with single-molecule spectroscopy. Proc Natl Acad Sci U S A 109:16155–16160. https://doi.org/10.1073/pnas.1207719109

Huang J, Mackerell AD (2013) CHARMM36 all-atom additive protein force field: Validation based on comparison to NMR data. J Comput Chem 34:2135–2145. https://doi.org/10.1002/jcc.23354

Humphrey W, Dalke A, Schulten K (1996) VMD: Visual Molecular Dynamics. J Mol Graph 14:33–38

Jha AK, Colubri A, Freed KF, Sosnick TR (2005) Statistical coil model of the unfolded state: Resolving the reconciliation problem. Proc Natl Acad Sci U S A 102:13099–13104. https://doi.org/10.1073/pnas.0506078102

Kenzaki H, Koga N, Hori N, et al (2011) CafeMol: A coarse-grained biomolecular simulator for simulating proteins at work. J Chem Theory Comput 7:1979–1989. https://doi.org/10.1021/ct2001045

Konovalova A, Kahne DE, Silhavy TJ (2017) Outer Membrane Biogenesis. Annu Rev Microbiol 71:539–556. https://doi.org/https://doi.org/10.1146/annurev-micro-090816-093754

Krainer G, Gracia P, Frotscher E, et al (2017) Slow Interconversion in a Heterogeneous Unfolded-State Ensemble of Outer-Membrane Phospholipase A. Biophys J 113:1280–1289. https://doi.org/10.1016/j.bpj.2017.05.037

Larsen AH, Wang Y, Bottaro S, et al (2020) RESEARCH ARTICLE Combining molecular dynamics simulations with small-angle X-ray and neutron scattering data to study multi-domain proteins in solution. PLoS Comput Biol 16:1–29. https://doi.org/10.1371/journal.pcbi.1007870

Marx DC, Plummer AM, Faustino AM, et al (2020) SurA is a cryptically grooved chaperone that expands unfolded outer membrane proteins. Proc Natl Acad Sci U S A 117:28026–28035. https://doi.org/10.1073/pnas.2008175117

Pelikan M, Hura GL, Hammel M (2009) Structure and flexibility within proteins as identified through small angle X-ray scattering. Gen Physiol Biophys 28:174–189. https://doi.org/10.4149/gpb_2009_02_174

Peran I, Holehouse AS, Carrico IS, et al (2019) Unfolded states under folding conditions accommodate sequence-specific conformational preferences with random coil-like dimensions. Proc Natl Acad Sci U S A 116:12301–12310. https://doi.org/10.1073/pnas.1818206116

Phillips JC, Braun R, Wang W, et al (2005) Scalable molecular dynamics with NAMD. J Comput Chem 26:1781–1802. https://doi.org/10.1002/jcc.20289

Philo JS (2006) Improved methods for fitting sedimentation coefficient distributions derived by time-derivative techniques. Anal Biochem 354:238–246. https://doi.org/10.1016/j.ab.2006.04.053

Plummer AM, Fleming KG (2016) From Chaperones to the Membrane with a BAM! Trends Biochem Sci 41:872–882. https://doi.org/10.1016/j.tibs.2016.06.005

Potrzebowski W, Trewhella J, Andre I (2018) Bayesian inference of protein conformational ensembles from limited structural data. PLoS Comput Biol 14:1–27. https://doi.org/10.1371/journal.pcbi.1006641

Riback JA, Bowman MA, Zmyslowski AM, et al (2017) Innovative scattering analysis shows that hydrophobic disordered proteins are expanded in water. Science (80-) 358:238–241

Rotkiewicz P, Skolnick J (2008) Fast Procedure for Reconstruction of Full-Atom Protein Models from Reduced Representations. J Comput Chem 29:1460–1465. https://doi.org/10.1002/jcc

Róycki B, Kim YC, Hummer G (2011) SAXS ensemble refinement of ESCRT-III CHMP3 conformational transitions. Structure 19:109–116. https://doi.org/10.1016/j.str.2010.10.006

Sherman E, Haran G (2006) Coil – globule transition in the denatured state of a small protein. Proc Natl Acad Sci U S A 103:11539–11543

Shevchuk R, Hub JS (2017) Bayesian refinement of protein structures and ensembles against SAXS data using molecular dynamics. PLoS Comput Biol 13:1–27. https://doi.org/10.1371/journal.pcbi.1005800

Shrestha UR, Juneja P, Zhang Q, et al (2019) Generation of the configurational ensemble of an intrinsically disordered protein from unbiased molecular dynamics simulation. Proc Natl Acad Sci U S A 116:20446–20452. https://doi.org/10.1073/pnas.1907251116

Tan AE, Burgess NK, DeAndrade DS, et al (2010) Self-association of unfolded outer membrane proteins. Macromol Biosci 10:763–767

Tesei G, Schulze TK, Crehuet R, Lindorff-Larsen K (2021) Accurate model of liquid-liquid phase behavior of intrinsically disordered proteins from optimization of single-chain properties. Proc Natl Acad Sci U S A 118:. https://doi.org/10.1073/pnas.2111696118

Tezuka-Kawakami T, Gell C, Brockwell DJ, et al (2006) Urea-Induced Unfolding of the Immunity Protein Im9 Monitored by spFRET. Biophys J L42–L44. https://doi.org/10.1529/biophysj.106.088344

Tomasek D, Kahne D (2021) The assembly of β-barrel outer membrane proteins. Curr Opin Microbiol 60:16–23. https://doi.org/10.1016/j.mib.2021.01.009

Ulrich T, Rapaport D (2015) Biogenesis of beta-barrel proteins in evolutionary context. Int J Med Microbiol 305:259–264. https://doi.org/10.1016/j.ijmm.2014.12.009

Zheng W, Borgia A, Buholzer K, et al (2016) Probing the Action of Chemical Denaturant on an Intrinsically Disordered Protein by Simulation and Experiment. J Am Chem Soc 138:11702–11713. https://doi.org/10.1021/jacs.6b05443. Probing PyMol Molecular Graphics System

