## Supplemental Information for "Generation of Unfolded Outer Membrane Protein Ensembles Targeted by Hydrodynamic Properties"

### Supplemental Methods

*Purification of OMPs from Inclusion Bodies*

Cloning of $OmpA_{171}$ into a pET11a vector with ampicillin resistance is described in (Danoff and Fleming 2011). Cloning of OmpX into a pET11a vector with ampicillin resistance is described in (Burgess et al. 2008). Glycerol stocks of plasmid transformed into HMS174(DE3) cells that had been stored at -80 °C were used to start 5 mL terrific broth (TB, Fisher) overnight cultures containing 100 ug/mL ampicillin (Sigma). The following day, overnight cultures were used to inoculate 500 mL of TB containing 100 ug/mL ampicillin, and cultures were grown at 37 °C with shaking until reaching an optical density at 600 nm ($OD_{600}$) of 1.0. Protein expression was induced by supplementing cultures with 1 mM isopropyl-β-D1-thiogalactopyranoside (IPTG) (ThermoScientific) and incubating them at 37 °C with shaking for an additional 4-6 hours. Cells were then harvested by centrifugation in a Beckman J2-MI centrifuge using a JA-10 rotor at 5000 rpm and 4 °C for 30 min. Cell pellets were stored at -20 C until lysis.

A pellet from a 500 mL growth was resuspended in 25 mL of OMP Lysis Buffer (50 mM Tris, 40 mM ethylenediaminetetraacetic acid (EDTA), pH 8) and subsequently lysed using an Avestin Emulsiflex homogenizer. After lysis, Brij-L23 (Sigma) was added to a final concentration of 0.1%, and full lysates were centrifuged (Beckman J2-MI centrifuge, JA-10 rotor) at 5000 rpm and 4 °C for 30 minutes. Pellets containing the isolated inclusion bodies were washed twice by resuspending in 25 mL of OMP Wash Buffer (10 mM Tris, 1 mM EDTA, pH 8) and subsequently centrifuging under the same conditions listed above. The purified inclusion bodies were resuspended a final time in 10 mL of OMP Wash Buffer and were aliquoted into 1 mL portions in 1.5 mL Eppendorf tubes before centrifuging a final time at 14,000 rpm and room temperature for 30 minutes using a table-top centrifuge (Eppendorf). The supernatant was discarded, and inclusion body pellets were stored at -20 °C.

An inclusion body aliquot was resuspended in 1 mL of either 20 mM Tris or 20 mM sodium phosphate buffer at pH 8 then pelleted at 14,000 rpm and room temperature for 5 minutes using a table-top centrifuge (Eppendorf). The inclusion body pellet was dissolved in 1.2 mL of 8 M Urea in the chosen buffer background (Tris or phosphate) by incubation at room temperature for >15 minutes. Once the pellet was fully dissolved, contaminating nucleic acids were removed by centrifugation at 14,000 rpm and room temperature for 5 minutes using a table-top centrifuge (Eppendorf). The supernatant was carefully pipetted off, and final sample purity and concentration were assessed by collecting a UV-visible absorbance wavelength spectrum. Samples were deemed pure of nucleic acid contaminants if the $A_{260}/A_{280} \leq 0.6$, and protein concentration was determined using the theoretical extinction coefficients calculated using the Edelhoch method (Edelhoch 1967) in SEDNTERP (Laue et al. 1992) ($\varepsilon_{280}$ of $OmpA_{171}$ = 45,090 $M^{-1}$ $cm^{-1}$ and $\varepsilon_{280}$ of OmpX = 31860 $M^{-1}$ $cm^{-1}$). Protein was diluted with 8 M Urea in the appropriate buffer (Tris or Phosphate) to a final concentration of approximately 80 μM and then aliquoted and stored at -80 °C until use.

### Supplemental Figures

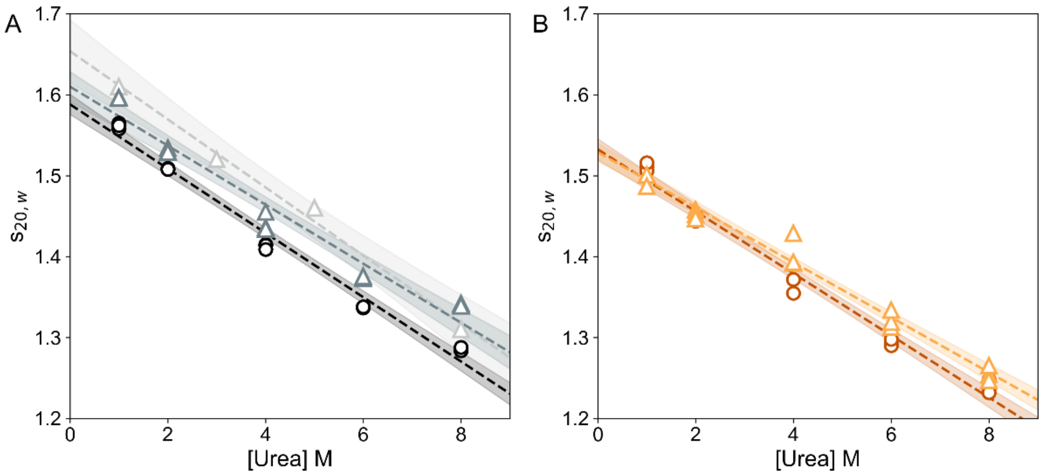

*Supplemental Fig 1. The buffer background negligibly affects the sedimentation of a uOMP ensemble.* For both uOMPs, the protein concentration was 2 μM in a buffer background of 20 mM sodium phosphate (dark color, open circles) or 20 mM Tris (light color, open triangles). Previously published data for uOmpA$_{171}$ in a Tris background is reproduced from Danoff and Fleming (Danoff and Fleming 2011) in the lightest grey color. In phosphate, $s_0$ = 1.59 (1.58-1.59) Svedbergs for (A) uOmpA$_{171}$ and $s_0$= 1.53 (1.52-1.54) Svedbergs for (B) uOmpX. In tris, $s_0$ = 1.61 (1.59-1.63) Svedbergs for (A) uOmpA$_{171}$ and $s_0$= 1.53 (1.52-1.54) Svedbergs for (B) uOmpX. Absorbance was measured at 230 nm while spinning at 50000 rpm and 25 °C. Data were analyzed using dc/dt+ to determine $s_{20,w}$ values. Each experiment was performed in triplicate, and all three data points at each urea concentration are included in the liner fit. Shaded regions represent the 95% confidence interval on the fit line.

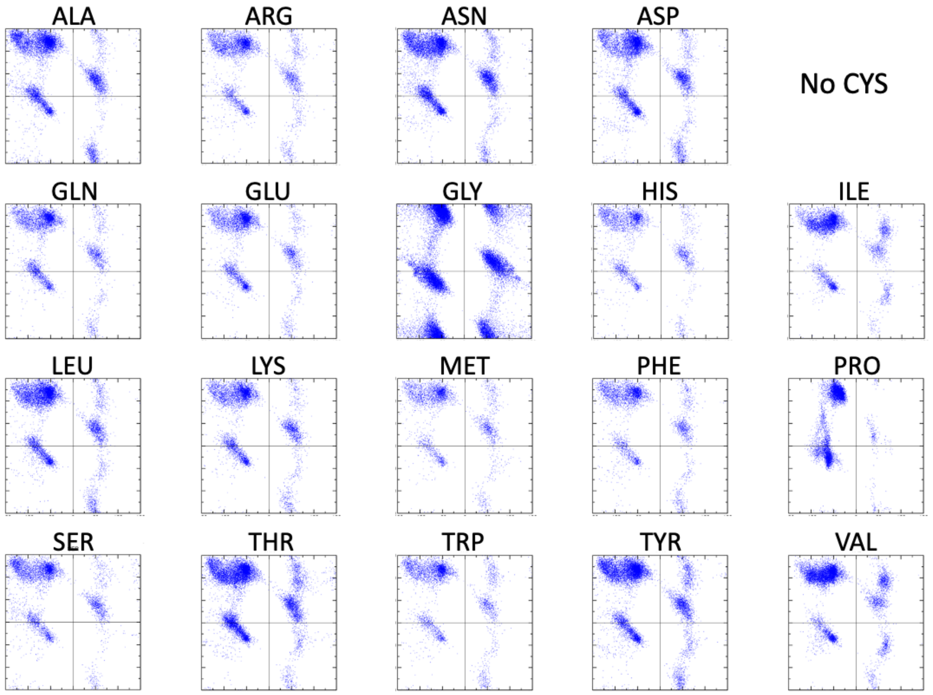

*Supplemental Fig 2. All residue types in refined OMP structures have favorable backbone dihedral angles.* Plots of ψ versus ϕ angles for an all-atom relaxed ensemble of 1000 uOMP structures. Each plot has axes of -180° to +180°. $uOmpA_{171}$ and uOmpX do not contain cysteine. The angle distributions are qualitatively consistent with ψ,ϕ angle distributions observed in an unbiased coil library (Beck et al. 2008).

### Supplemental Tables

*Supplemental Table 1. Summary of experimental SV-AUC values.* All experiments were performed at 25 °C. All densities (ρ), viscosities (η), and partial specific volumes (**$\bar{v}$**) reported here assume a temperature of 25 °C.

| **Protein** | **Buffer** | **[Urea]** (M) | **ρ** (g/mL) | **η** (cP) | **$\bar{v}$** (mL/g) | <s> Svedbergs | <$s_{20,w}$> Svedbergs | **Fit $s_{20,w}$** Svedbergs | **MW** (kDa) |
|---|---|---|---|---|---|---|---|---|---|
| uOmpA | Phosphate | 1 | 1.01527 | 0.93316 | 0.7229 | 1.516 | 1.487 | 1.558 | 18.71 |
| | | | | | | 1.525 | 1.498 | 1.565 | 18.81 |
| | | | | | | 1.518 | 1.489 | 1.562 | 18.65 |
| | | 2 | 1.03108 | 0.97648 | 0.7217 | 1.355 | 1.452 | 1.509 | 19.87 |
| | | | | | | 1.351 | 1.450 | 1.508 | 19.76 |
| | | | | | | 1.356 | 1.454 | 1.508 | 19.36 |
| | | 4 | 1.06210 | 1.0856 | 0.7193 | 1.045 | 1.362 | 1.415 | 20.34 |
| | | | | | | 1.043 | 1.359 | 1.414 | 20.45 |
| | | | | | | 1.042 | 1.359 | 1.409 | 20.12 |
| | | 6 | 1.09147 | 1.2459 | 0.7169 | 0.766 | 1.256 | 1.337 | 20.20 |
| | | | | | | 0.765 | 1.252 | 1.338 | 20.56 |
| | | | | | | 0.768 | 1.259 | 1.338 | 20.00 |
| | | 8 | 1.12146 | 1.4854 | 0.7145 | 0.557 | 1.200 | 1.287 | 20.45 |
| | | | | | | 0.555 | 1.196 | 1.284 | 20.59 |
| | | | | | | 0.558 | 1.203 | 1.288 | 20.59 |
| | Tris | 1 | 1.01299 | 0.92900 | 0.7229 | 1.565 | 1.520 | 1.596 | 18.85 |
| | | | | | | 1.557 | 1.512 | 1.597 | 19.21 |
| | | | | | | 1.552 | 1.506 | 1.596 | 18.77 |
| | | 2 | 1.02881 | 0.97232 | 0.7217 | 1.399 | 1.487 | 1.534 | 20.26 |
| | | | | | | 1.376 | 1.461 | 1.530 | 20.06 |
| | | | | | | 1.421 | 1.508 | 1.529 | 20.19 |
| | | 4 | 1.05982 | 1.0815 | 0.7193 | 1.062 | 1.371 | 1.436 | 21.07 |
| | | | | | | 1.055 | 1.360 | 1.434 | 20.80 |
| | | | | | | 1.078 | 1.390 | 1.455 | 20.90 |
| | | 6 | 1.08919 | 1.2418 | 0.7168 | 0.795 | 1.286 | 1.373 | 21.54 |
| | | | | | | 0.793 | 1.284 | 1.376 | 21.53 |
| | | | | | | 0.801 | 1.297 | 1.375 | 21.01 |
| | | 8 | 1.11936 | 1.4813 | 0.7144 | 0.590 | 1.258 | 1.342 | 20.84 |
| | | | | | | 0.581 | 1.238 | 1.341 | 20.62 |
| | | | | | | 0.586 | 1.250 | 1.339 | 21.01 |
| uOmpX | Phosphate | 1 | 1.01527 | 0.93316 | 0.7155 | 1.522 | 1.491 | 1.512 | 15.21 |
| | | | | | | 1.508 | 1.477 | 1.506 | 15.24 |
| | | | | | | 1.535 | 1.504 | 1.516 | 15.14 |
| | | 2 | 1.03108 | 0.97648 | 0.7142 | 1.340 | 1.432 | 1.448 | 16.21 |
| | | | | | | 1.322 | 1.414 | 1.444 | 16.07 |
| | | | | | | 1.329 | 1.420 | 1.451 | 16.18 |
| | | 4 | 1.06210 | 1.0856 | 0.7116 | 1.016 | 1.315 | 1.355 | 16.44 |
| | | | | | | 1.016 | 1.316 | 1.372 | 17.08 |
| | | | | | | 1.021 | 1.322 | 1.372 | 17.05 |
| | | 6 | 1.09147 | 1.2459 | 0.7090 | 0.752 | 1.216 | 1.292 | 17.36 |
| | | | | | | 0.754 | 1.221 | 1.290 | 16.90 |
| | | | | | | 0.757 | 1.224 | 1.298 | 17.12 |
| | | 8 | 1.12146 | 1.4854 | 0.7064 | 0.547 | 1.159 | 1.248 | 17.50 |
| | | | | | | 0.538 | 1.141 | 1.238 | 17.24 |
| | | | | | | 0.532 | 1.126 | 1.232 | 16.83 |
| | Tris | 1 | 1.01299 | 0.92900 | 0.7155 | 1.463 | 1.420 | 1.501 | 16.50 |

| | | | | | | | | | |
|---|---|---|---|---|---|---|---|---|---|
| | | | | | | 1.445 | 1.401 | 1.501 | 16.28 |
| | | | | | | 1.424 | 1.381 | 1.487 | 16.51 |
| | | 2 | 1.02881 | 0.97232 | 0.7142 | 1.302 | 1.375 | 1.458 | 17.37 |
| | | | | | | 1.301 | 1.373 | 1.452 | 17.01 |
| | | | | | | 1.276 | 1.353 | 1.447 | 16.99 |
| | | 4 | 1.05982 | 1.0815 | 0.7115 | 1.054 | 1.352 | 1.429 | 15.62 |
| | | | | | | 1.014 | 1.302 | 1.392 | 17.83 |
| | | | | | | 1.013 | 1.298 | 1.393 | 17.53 |
| | | 6 | 1.08919 | 1.2418 | 0.7088 | 0.755 | 1.210 | 1.313 | 17.62 |
| | | | | | | 0.745 | 1.195 | 1.319 | 17.89 |
| | | | | | | 0.760 | 1.217 | 1.334 | 17.10 |
| | | 8 | 1.11936 | 1.4813 | 0.7062 | 0.554 | 1.161 | 1.254 | 18.02 |
| | | | | | | 0.544 | 1.140 | 1.247 | 17.70 |
| | | | | | | 0.559 | 1.171 | 1.265 | 17.82 |

**Supplemental References**

Beck DAC, Alonso DOV, Inoyama D, Daggett V (2008) The intrinsic conformational propensities of the 20 naturally occurring amino acids and reflection of these propensities in proteins. Proc Natl Acad Sci U S A 105:12259–12264. https://doi.org/10.1073/pnas.0706527105

Burgess NK, Dao TP, Stanley AM, Fleming KG (2008) β-Barrel proteins that reside in the Escherichia coli outer membrane in vivo demonstrate varied folding behavior in vitro. J Biol Chem 283:26748–26758. https://doi.org/10.1074/jbc.M802754200

Danoff EJ, Fleming KG (2011) The soluble, periplasmic domain of OmpA folds as an independent unit and displays chaperone activity by reducing the self-association propensity of the unfolded OmpA transmembrane β-barrel. Biophys Chem 159:194–204. https://doi.org/10.1016/j.bpc.2011.06.013

Edelhoch H (1967) Spectroscopic Determination of Tryptophan and Tyrosine in Proteins. Biochemistry 6:1948–1954. https://doi.org/https://doi.org/10.1021/bi00859a010

Laue TM, Shah BD, Ridgeway TM, Pelletier SL (1992) Computer-aided interpretation of analytical sedimentation data for proteins. In: Harding S, Rowe A, Hoarton J (eds) Analytical Ultracentrifugation in Biochemistry and Polymer Science. Royal Society of Chemistry, Cambridge, UK, pp 90–125
